# No evidence for changes in GABA concentration, functional connectivity, or working memory following continuous theta burst stimulation over dorsolateral prefrontal cortex

**DOI:** 10.1101/2021.06.17.448448

**Authors:** Tribikram Thapa, Joshua Hendrikse, Sarah Thompson, Chao Suo, Mana Biabani, James Morrow, Kate E Hoy, Paul B Fitzgerald, Alex Fornito, Nigel C Rogasch

## Abstract

Continuous theta burst stimulation (cTBS) is thought to reduce cortical excitability and modulate functional connectivity, possibly by altering cortical inhibition at the site of stimulation. However, most evidence comes from the motor cortex and it remains unclear whether similar effects occur following stimulation over other brain regions. We assessed whether cTBS over left dorsolateral prefrontal cortex altered gamma aminobutyric acid (GABA) concentration, functional connectivity and brain dynamics at rest, and brain activation and memory performance during a working memory task. Seventeen healthy individuals participated in a randomised, sham-controlled, cross-over experiment. Before and after either real or sham cTBS, magnetic resonance spectroscopy was obtained at rest to measure GABA concentrations. Functional magnetic resonance imaging (fMRI) was also recorded at rest and during an n-back working memory task to measure functional connectivity, regional brain activity (low-frequency fluctuations), and task-related patterns of brain activity. We could not find evidence for changes in GABA concentration (*P*=0.66, Bayes factor [BF_10_]=0.07), resting-state functional connectivity (*P*_(FWE)_>0.05), resting-state low-frequency fluctuations (*P*=0.88, BF_10_=0.04), blood-oxygen level dependent activity during the n-back task (*P*_(FWE)_ >0.05), or working memory performance (*P*=0.13, BF_10_=0.05) following real or sham cTBS. Our findings add to a growing body of literature suggesting the effects of cTBS are highly variable between individuals and question the notion that cTBS is a universal ‘inhibitory’ paradigm.

## 1. Introduction

Continuous theta burst stimulation (cTBS) is a form of patterned repetitive transcranial magnetic stimulation (TMS) capable of inducing plasticity in neural circuits and influencing cognition (Hallett, 2007). In humans, cTBS involves applying triplets of pulses at 50 Hz (i.e., a gamma rhythm) repeated at 5 Hz intervals (i.e., a theta rhythm) typically for 600 pulses (Huang et al., 2005). When applied over the motor cortex, early studies observed a reduction in the amplitude of motor-evoked potentials (MEPs) lasting ∼30-60 mins (Di Lazzaro et al., 2005; Huang et al., 2005; Huang et al., 2007). These changes were blocked by pharmacological antagonists of n-methyl-d-aspartate (NMDA) receptors (Huang et al., 2005) and voltage-gated calcium channels (Wankerl et al., 2010), suggesting that cTBS results in a transient reduction in cortical excitability possibly reflecting LTD-like mechanisms. In addition to measures of excitability, cTBS over motor cortex reduces paired-pulse TMS measures of cortical inhibition (Huang et al., 2005; Murakami et al., 2008; McAllister et al., 2009; Goldsworthy et al., 2013) and increases concentrations of the inhibitory neurotransmitter gamma-amino butyric acid (GABA) measured with magnetic resonance spectroscopy (MRS) (Stagg et al., 2009), suggesting an interaction with local inhibitory circuits. Outside of the stimulated region, cTBS increases MEP amplitude in the contralateral motor cortex (Suppa et al., 2008) and alters functional connectivity measured using functional magnetic resonance imaging (fMRI) between the stimulated motor region and other cortical networks (Cocchi et al., 2015; Steel et al., 2016), suggesting that cTBS has broader effects on brain activity beyond the site of stimulation. Together, these findings from motor cortex have led to the view that cTBS is an ‘inhibitory’ repetitive TMS paradigm which decreases excitability at the site of stimulation, possibly by altering local cortical inhibition (Suppa et al., 2016). The local changes in cortical excitability and inhibition are then thought to impact connectivity with downstream cortical sites, resulting in a rebalancing of large-scale cortical networks at rest and during tasks (Sale et al., 2015).

The concept of cTBS as an ‘inhibitory’ paradigm has been widely adopted in attempts to study the causal role of brain regions in cognitive ability (Ngetich et al., 2020). For example, neuroimaging (Rottschy et al., 2012) and lesion studies (Barbey et al., 2013) have implicated the dorsolateral prefrontal cortex (DLPFC) as an important region for working memory. Subsequently, several studies have attempted to reduce the activity/connectivity of DLPFC with cTBS in healthy individuals and then test the impact on working memory performance. While two studies have reported reduced working memory performance following cTBS over DLPFC (Schicktanz et al., 2015; Vékony et al., 2018), this result has not always been replicated (Viejo-Sobera et al., 2017). Furthermore, it remains largely unclear from these studies how cTBS over DLPFC altered brain activity/connectivity leading to changes in working memory performance.

Despite the widespread assumption that cTBS is a universal ‘inhibitory’ rTMS paradigm regardless of the region stimulated, this notion has been challenged on several fronts. First, there is a growing body of evidence suggesting that the outcomes of cTBS may vary across the cortex. For example, Cocchi et al. (2016) found that cTBS over frontal eye fields reduced resting-state functional connectivity with the stimulated site, whereas cTBS over occipital cortex increased connectivity. Computational modelling suggested that differences in local low-frequency fluctuations between sites could have explained the difference in connectivity outcome. Other ‘inhibitory’ rTMS paradigms (e.g., 1 Hz rTMS) have also resulted in similar divergent changes in connectivity when targeting different regions (Castrillon et al., 2020), suggesting the directionality of rTMS outcomes derived from motor cortex may not necessarily translate to other stimulated brain regions or networks. Second, the outcomes of cTBS may also heavily depend on the individual. For example, several studies have found that MEPs are reduced in <50% of individuals following cTBS using the ‘standard’ parameters (600 pulses, subthreshold stimulation intensity etc.) (Goldsworthy et al., 2012; Hamada et al., 2013). Indeed, recent meta-analyses have found strong evidence for publication bias within the cTBS literature (Chung et al., 2016), suggesting potential under-reporting of inter-individual variability following cTBS. Together, these findings have led to calls for careful evaluation of how rTMS paradigms like cTBS alter brain function when targeting non-motor regions, especially when trying to establish causality between neuronal and behavioural effects (Bergmann & Hartwigsen, 2021).

The aim of this study was to perform a comprehensive, multi-modal assessment of how cTBS over DLPFC alters brain activity and connectivity, as well as subsequent working memory performance. We measured changes in local and remote GABA concentration, functional connectivity and low-frequency fluctuations at rest, blood oxygenation level-dependent (BOLD) activity during an n-back working memory task and working memory performance. Based on available evidence, we expected increases in local GABA concentration, reductions in low-frequency fluctuations, reductions in functional connectivity with the stimulated site, reductions in DLPFC BOLD activity during working memory, and impaired working memory performance following cTBS. Where possible, we used a combination of frequentist and Bayesian statistical tests to assess evidence for and against the null-hypotheses.

## 2. Materials and Methods

### 2.1 Ethics approval

This study was approved by the Monash University Human Research Ethics Committee (Project ID: 6054) and performed in accordance with the standards set by the Declaration of Helsinki.

### 2.2 Participants

Eighteen healthy, right-handed individuals were recruited for the study. One participant was excluded due to missing data, therefore data is presented from 17 participants (8 males, 9 females; age: 32.53 ± 8.64 years). All participants met inclusion/exclusion criteria for MRI and TMS. Participants were reimbursed $90 AUD upon study completion.

### 2.3 Experimental protocol

All participants attended two experimental sessions at least 7 days apart (fig. 1). In each session, participants completed a baseline MRI scan. Participants then moved to a room adjacent to the scanner, where resting motor threshold (RMT) was obtained over the left primary motor cortex, followed by cTBS to the left DLPFC (real or sham). Participants returned to the scanner immediately following stimulation for a post-cTBS MRI scan. The order of real or sham cTBS across sessions was counterbalanced between individuals.

**Figure 1:**
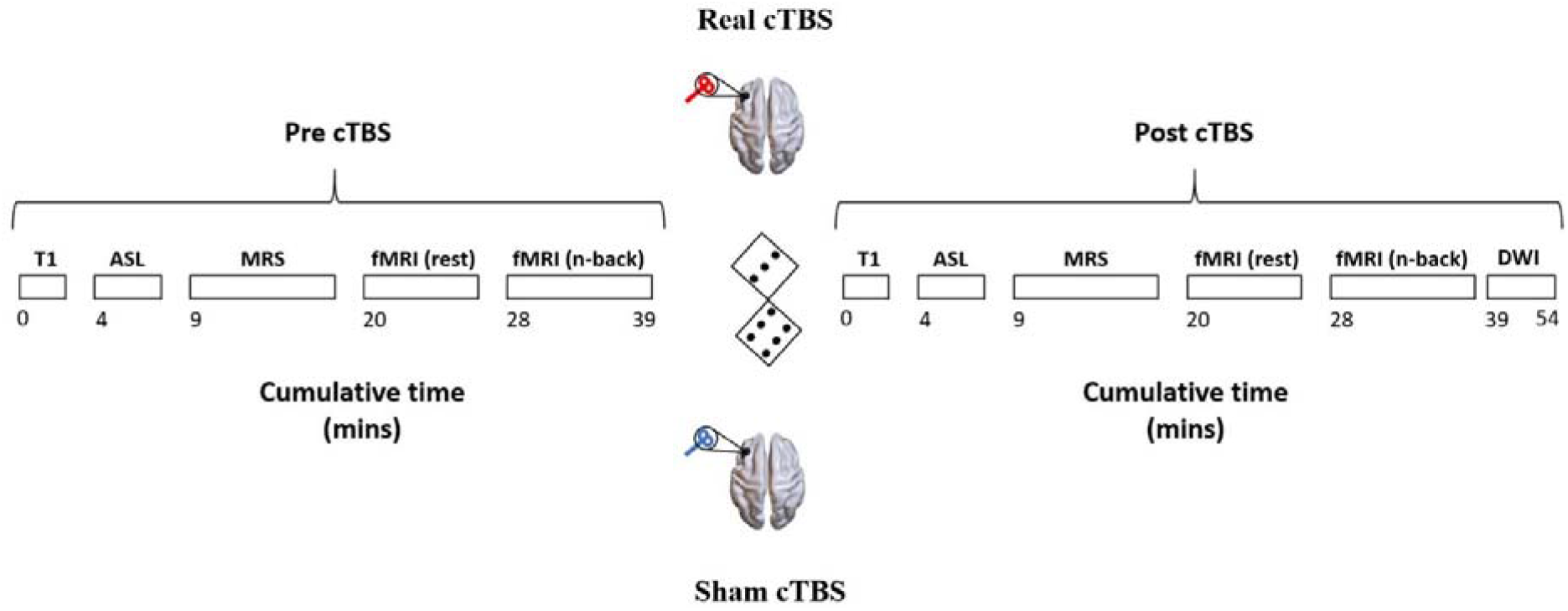
Diagram showing the magnetic resonance imaging (MRI) sequences collected before and after continuous theta burst stimulation (cTBS) in each experimental session. Cumulative time refers to the time elapsed since the beginning of the scan (separate for pre and post scans). ASL, arterial spin labelling; MRS, magnetic resonance spectroscopy; fMRI, functional magnetic resonance imaging; DWI, diffusion weighted imaging.

### 2.4 Data Acquisition

#### 2.4.1 Magnetic resonance imaging (MRI)

A Siemens Biograph mMR scanner with a 32-channel head coil was used to scan all participants. Identical scans were acquired pre- and post-cTBS. First, T1-weighted (T1w) structural images were acquired (Magnetization Prepared Rapid Gradient Echo, TA: 3.49 mins, TR: 1640 ms, TE: 2.34 ms, Voxel size: 1 mm^3^, Flip angle: 8□, Slices: 176). While voxel placement for MRS was taking place, participants underwent an arterial spin labelling scan (data not analysed). Spectroscopy data was then collected from the left DLPFC (25 × 25 × 13 mm; stimulation site) and visual cortex (13 × 25 × 25 mm; control site) using a GABA-edited Mescher-Garwood Point Resolved Spectroscopy (MEGA-PRESS) sequence (96 ON-OFF averages, TE=68ms, TR=1500ms, edit pulse at 1.9ppm with 46Hz bandwidth). To place the DLPFC voxel, the MNI coordinate [-39 30 21] was defined in standard space as a reference point. Radiographers later used this information and the three orthogonal views of T1w images to further identify the DLPFC region in subject specific space and placed the voxel carefully to avoid the skull and cerebrospinal fluid regions. The visual cortex voxel was located over the left primary visual cortex. Both regions of interest are illustrated in Figure 2. For each region, unsuppressed water was also obtained using the same parameters (except 8 ON-OFF averages). This was followed by EPI sequences acquired at rest with eyes open (TA: 7.52 mins, TR: 1368 ms, TE: 67.2 ms, Voxel size: 3.2 × 3.2 × 3.0 mm Flip angle: 50□, Vol: 340) and during two runs of an n-back task (TA: 5.07 mins for each run, TR: 2500 ms, TE: 30 ms, Voxel size: 3 mm^3^, Flip angle: 90□, Vol: 119). The MEGA-PRESS sequence was deliberately acquired before EPI acquisition to minimise signal drift caused by heating of the coil assembly in EPI sequences (Harris et al., 2014; Tsai et al., 2016). At the end of the post-cTBS session, a diffusion-weighted scan was also obtained (data not analysed).

**Figure 2:**
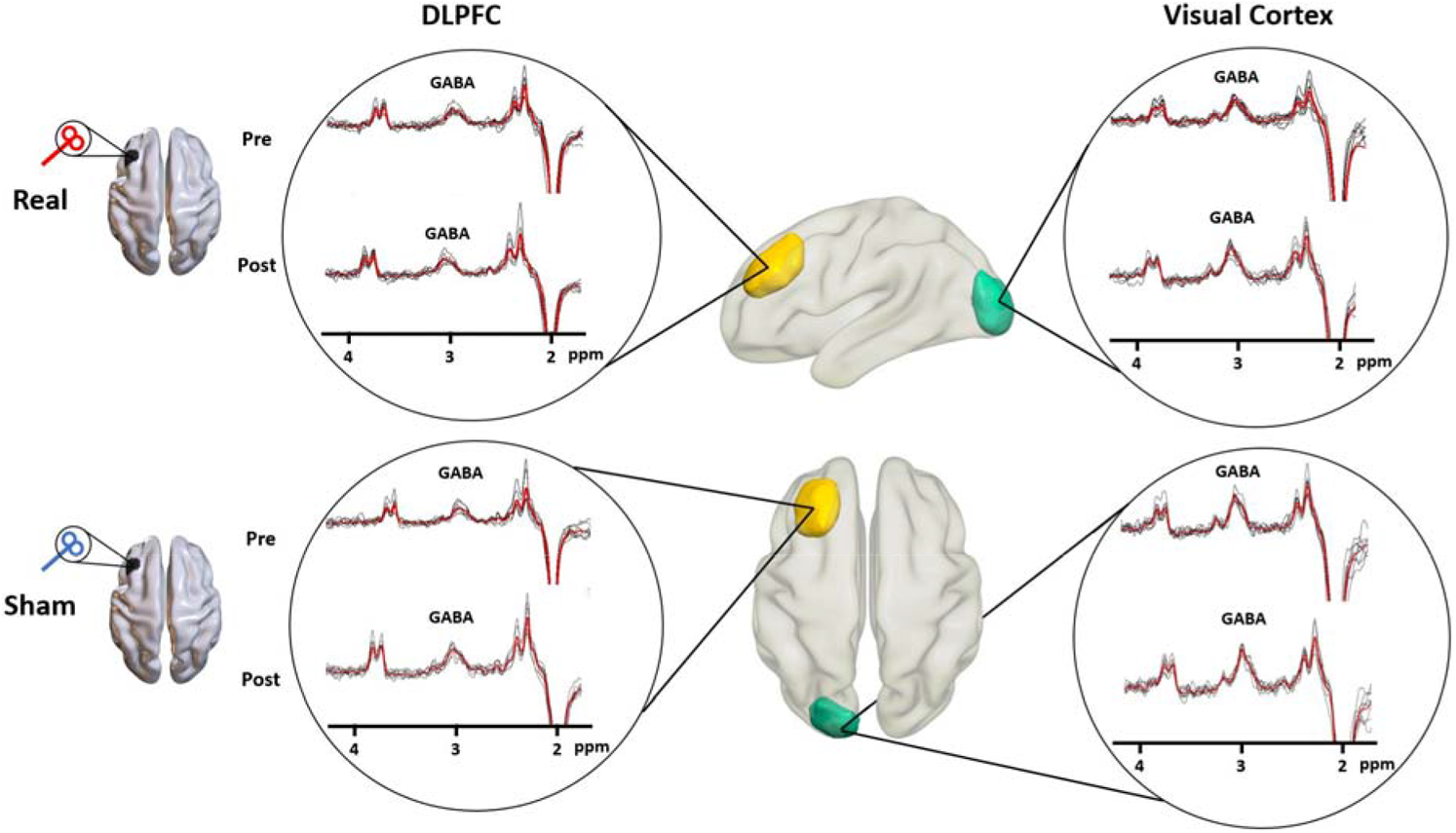
Gamma aminobutyric acid (GABA) concentrations measured using magnetic resonance spectroscopy before and after continuous theta burst stimulation (cTBS). Individual spectra with the GABA peak are shown in black traces for each condition (real and sham cTBS) and time-point (pre and post) within each circle. Red traces represent the group average (n = 8). Mean voxels from which GABA concentrations were calculated are also shown from the dorsolateral prefrontal cortex (DLPFC, yellow) and visual cortex (green). Voxels were reconstructed using each subject’s T1 and spectroscopy file. Individual T1s were aligned using 6-degrees of freedom to the standard MNI template along with the reconstructed voxel of interest. The mean voxel was generated by overlapping all aligned individual voxel of interest with a threshold of 0.2.

#### 2.4.2 N-back task

Participants performed the n-back task across two runs at each time-point (Harding et al., 2016). Each run comprised of 6 blocks alternating between 0-back and 2-back conditions (3 blocks of each) with 14 trials in each block. Task stimuli consisting of individual letters were presented for 500 ms and separated by a pseudorandomly varying interstimulus interval of 1250 ms, 1500 ms, or 1750 ms. The first run began with a 0-back condition, whereas the second began with a 2-back condition. For the 0-back condition, participants were instructed to press a button with their right hand every time a target letter (B) appeared on the screen. For the 2-back condition, participants were instructed to press a button only when the current letter matched a letter presented two trials earlier. There were 4 target trials in each block of 14 stimuli. A fixation cross was presented before and between each block.

#### 2.4.3 Transcranial magnetic stimulation (TMS)

Transcranial magnetic stimulation (TMS) was delivered using a B-65 figure-of-eight cooled coil (75 mm diameter) attached to a MagVenture MagPro X100 stimulator. Biphasic pulses were given with the coil handle positioned approximately 90° to the targeted gyrus such that an anterior-posterior followed by posterior-anterior current flow was achieved in the underlying cortex. A stereotaxic infrared neuronavigation system (Brainsight, Rogue Research) was used to monitor TMS coil placement, and to personalise TMS target sites on each subject’s T1 anatomical image. RMT was determined from the hotspot over left motor cortex corresponding to the right first dorsal interosseous (FDI) muscle and was defined as the lowest stimulation intensity required to elicit a motor evoked potential >0.05 mV peak-to-peak amplitude in at least 5 out of 10 trials. Real cTBS was administered to the left DLPFC (MNI co-ordinates: −38, 30, 30) at 70% RMT in triplets of 50 Hz repeated every 5 Hz for 600 pulses. The stimulation site was chosen to target a common DLPFC region activated during working memory based on a meta-analysis of neuroimaging studies (Yarkoni et al., 2011). Sham cTBS was administered using the same parameters but with the coil held at a 90□ angle to the DLPFC so that the magnetic field ran perpendicular to scalp. Stimulation intensity for real and sham cTBS was adjusted for differences in scalp-to-cortex distance between the FDI hotspot and DLPFC using the formula suggested by Stokes and colleagues (Stokes et al., 2005).

### 2.5. Data analysis

#### 2.5.1 Magnetic resonance spectroscopy

Gannet (version 3.1) was used to analyse MEGA-PRESS spectra (Edden et al., 2014). Estimates of GABA concentration were calculated from voxels located at the DLPFC and visual cortex relative to the unsuppressed water signal. Here, raw Siemens ‘twix’ files of both water-suppressed and non-suppressed scans were first preprocessed using Gannet. After standard Gannet preprocessing, a GABA peak at 3.0 ppm was fitted using the GannetFit module, which further computed GABA concentration (in institutional units) relative to the unsuppressed water signal (Edden et al., 2014). Partial volume segmentation of T1-weighted anatomical images was conducted using Gannet to calculate the percentage of gray and white matter within the voxel of interest (GM% and WM%). These percentages were subsequently used to determine partial volume corrected GABA concentration [GABA_corrected=GABA/(GM%+WM%)] for the DLPFC and visual cortex. As GABA concentration estimates are highly susceptible to noise, the full width at half maximum (FWHM) of GABA peak and GABA peak fit error (relative to the unsuppressed water peak) were obtained from the GannetFit output and used for quality control. Spectra with an FWHM value of > 10Hz (suggesting a tendency to underestimate GABA (Provencher, 1993; Deelchand et al., 2015; Deelchand et al., 2018)), or a negative value for area (thought to reflect suboptimal water suppression and subsequently uncertain metabolite signal phase correction) were removed from the analysis (Alger, 2010).

#### 2.5.2 fMRI data analysis

fMRI and T1-weighted data were processed using fMRIPrep software (Esteban et al., 2019), as detailed in the following sections (https://fmriprep.org/en/stable/).

##### 2.5.2.1 Anatomical data preprocessing

Each participant’s T1w image was corrected for intensity non-uniformity (INU) with N4BiasFieldCorrection (Tustison et al., 2010), distributed with ANTs 2.2.0 (Avants et al., 2008), and used as T1w-reference throughout the workflow. The T1w-reference was then skull-stripped with a Nipype implementation of the antsBrainExtraction.sh workflow (from ANTs), using OASIS30ANTs as the target template. Brain tissue segmentation of cerebrospinal fluid (CSF), white-matter (WM) and gray-matter (GM) was performed on the brain-extracted T1w using fast (FSL 5.0.9) (Zhang et al., 2001). Brain surfaces were reconstructed using recon-all (FreeSurfer 6.0.1) (Dale et al., 1999), and the brain mask estimated previously was refined with a custom variation of the method to reconcile ANTs-derived and FreeSurfer-derived segmentations of the cortical gray-matter of Mindboggle (Klein et al., 2017). Volume-based spatial normalization to MNI152NLin2009cAsym standard space was performed through nonlinear registration with antsRegistration (ANTs 2.2.0), using brain-extracted versions of both T1w reference and the T1w template. ICBM 152 Nonlinear Asymmetrical template version 2009c was selected for spatial normalization:(Fonov et al., 2011) [TemplateFlow ID: MNI152NLin2009cAsym].

##### 2.5.2.2 Functional data preprocessing

The following preprocessing steps were performed for each of the BOLD runs (resting and task). First, a reference volume and its skull-stripped version was generated using a custom methodology of fMRIPrep. A deformation field to correct for susceptibility distortions was estimated based on fMRIPrep’s fieldmap-less approach. The deformation field is that resulting from co-registering the BOLD reference to the same-subject’s T1w-reference with its intensity inverted (Huntenburg, 2014; Wang et al., 2017). Registration was performed with antsRegistration (ANTs 2.2.0), and the process was regularised by constraining deformation to be nonzero only along the phase-encoding direction, and modulated with an average fieldmap template (Treiber et al., 2016). Based on the estimated susceptibility distortion, an unwarped BOLD reference was calculated for a more accurate co-registration with the anatomical reference. The BOLD reference was then co-registered to the T1w reference using bbregister (FreeSurfer) which implements boundary-based registration (Greve & Fischl, 2009). Co-registration was configured with nine degrees of freedom to account for distortions remaining in the BOLD reference. Head-motion parameters with respect to the BOLD reference (transformation matrices, and six corresponding rotation and translation parameters) were estimated before any spatiotemporal filtering using mcflirt (FSL 5.0.9) (Jenkinson et al., 2002). The BOLD time-series (including slice-timing correction when applied) were resampled onto their original, native, and MNI152NLin2009cAsym standard space by applying a single, composite transform to correct for head-motion and susceptibility distortions. These resampled BOLD time-series are preprocessed BOLD images. Additionally, automatic removal of motion artifacts using independent component analysis (ICA-AROMA) (Pruim et al., 2015) was performed on the preprocessed BOLD on MNI space time-series after removal of non-steady state volumes and spatial smoothing with an isotropic, Gaussian kernel of 6mm FWHM. Corresponding denoised runs were produced after such smoothing. Noise-regressors were then collected and placed in the corresponding confounds file. The three tissue-averaged signals were extracted from the CSF, the WM, and the whole-brain masks. These signals were then removed from the ICA-AROMA-denoised data using linear regression. A 6 mm smoothing kernel (SPM 12) was then applied to the denoised data.

Following preprocessing, resting-state functional connectivity was assessed in two ways: (1) a seed-based approach; and (2) a whole brain connectome approach. While the seed-based approach tests for changes in the targeted network, whole brain approach tests for changes in both targeted and non-targeted networks. For the seed-based analysis, a representative time course was extracted from a 5-mm spherical seed region placed in the DLPFC region and was used for statistical comparisons with time series from all other voxels. For the whole-brain connectome analysis, the brain was parcellated into 300 nodes using Schaefer’s parcellation (Schaefer et al., 2018) and the average time series was calculated across voxels within each node. Pearson correlations were then calculated between every pair of regional time courses.

As computational modeling has suggested changes in functional connectivity following cTBS may depend on low-frequency fluctuations at the target site (Cocchi et al., 2016), we also measured the fractional amplitude of low-frequency fluctuations (fALFF) from the DLPFC. fALFF was calculated on the time series extracted from the 5 mm spherical seed placed in the DLFPC using the *SP_fALFF* function from the highly comparative time series analysis toolbox (https://github.com/benfulcher/hctsa) (Fulcher et al., 2013; Fulcher & Jones, 2017).

#### 2.5.3 N-back performance

Working memory performance on the 2-back task was quantified using *d* Prime (d’) scores. d’ was calculated using the formula:

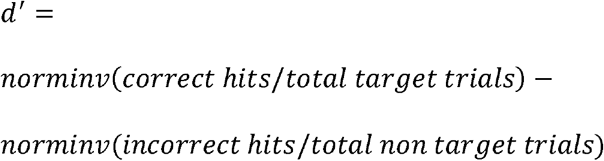

Here, norminv represents the inverse of the standard normal cumulative distribution, correct hits represents the number of target trials with a corresponding button press, and incorrect hits represents the number of non-target trials with a corresponding button press (i.e., a false alarm) (Haatveit et al., 2010). Mean reaction times (RT) to correct hits were also calculated. One subject was removed due to software malfunction during data acquisition. Therefore, working memory performance analysis was conducted on 16 subjects.

### 2.6 Statistical Analysis

#### 2.6.1 GABA concentrations, low frequency fluctuations, and working memory performance

A 2×2×2 repeated measures analysis of variance (ANOVA; SigmaPlot 12.3) was used to test for changes in GABA concentration following cTBS with main effects of TIME (pre and post), CONDITION (real and sham) and SITE (DLPFC and visual cortex). A 2×2 repeated measures ANOVA with factors TIME (pre and post) and CONDITION (real and sham) was used to assess changes in fALFF values and changes in working memory performance (d’ scores and RT) following cTBS. In all tests, a value of *P*<0.05 was considered statistically significant. When present, significant main effects and interactions were further explored using post-hoc t-tests.

To test the relative evidence for the alternative versus null hypothesis, the Bayesian equivalent of a repeated measures ANOVA was also conducted on GABA concentration, fALFF values, d’ scores and RT (JASP 0.10.2.0). A Bayes Factor (BF_10_) was calculated for main effects and interactions, which is a ratio that compares the likelihood of the data fitting the alternative hypothesis over the likelihood of the data fitting the null hypothesis (Goodman, 1999; Wagenmakers et al., 2018a; Wagenmakers et al., 2018b). A BF_10_>3 was taken as strong evidence supporting the alternative hypothesis, and a BF_10_<0.33 as strong evidence supporting the null (Wagenmakers et al., 2018a). The Cauchy parameter was set to a conservative default of 0.707 (Rouder et al., 2009; Ly et al., 2016).

#### 2.6.2 Resting state functional connectivity

For seed-based connectivity, data were analysed across two levels using SPM12 (https://www.fil.ion.ucl.ac.uk/spm/software/spm12/). At the 1^st^ level, four DLPFC functional connectivity contrast images (pre, post, pre-post and post-pre) were generated for the real and sham cTBS conditions separately for each individual. For the 2^nd^ level, two designs were used to assess group level differences. First, a flexible full factorial design using pre and post contrast images from the 1^st^ level was run to explore main effects (GROUP: real vs sham cTBS; TIME: pre vs post) and interaction effects at the group level. Second, a one sample t-test was run using all difference contrast images from the 1^st^ level. To correct for multiple comparison, a cluster level FWE correction with an initial threshold at *P*<0.001, k>10, and *P*_(FWE)_ <0.05 was considered significant.

Whole-brain functional connectivity was investigated using network-based statistics (NBS), and involved fitting a general linear model to each of 44,700 edges in the 300-by-300 region functional connectivity matrix (Zalesky et al., 2010). The GLM model was constructed including main factors of SUBJECT, TIME (pre and post) and GROUP (real and sham cTBS), as well as interaction of TIME by GROUP (4 levels). The contrasts of interest were set to examine any significant effect of TIME by GROUP interaction. NBS performs inference at the level of connected sets of supra-threshold edges (termed connected components), providing greater statistical power than mass univariate inference (Zalesky et al., 2010). We evaluated results with primary component-forming thresholds of T = 1, 1.5 and 2, and declared results significant if they passed a familywise error corrected threshold of 0.05. 1000 permutations were used for statistical inference (Zalesky et al., 2010).

#### 2.6.3 BOLD during the n-back task

Preprocessed data recorded during the n-back task were used to assess changes in task-related BOLD activity following cTBS. At the 1^st^ level, task parameters (duration and onset) corresponding to the 0-back and 2-back task conditions were coded as individual regressors alongside additional regressors for head motion (six degrees of freedom). The difference contrast between the 0-back greater than 2-back (0-back), and 2-back greater than 0-back (2-back) conditions were built at the 1^st^ level to represent brain deactivation and activation during n-back task performance, respectively. These contrasts were fed separately into the 2^nd^ level for group level analysis. At the 2^nd^ level, a flexible full factorial design was used to explore the main effects of CONDITION (real and sham cTBS) and TIME (pre and post), and interaction effects. A one-sample t-test using all difference contrast images from the 1^st^ level was also used to analyse differences in n-back task performance between real and sham cTBS at each time-point. To correct for multiple comparison, a cluster level FWE correction with a default initial threshold at *P*<0.001, k>10, and *P*_(FWE)_<0.05 was considered significant.

## 3. Results

### 3.1 Resting motor threshold

The mean (± SD) resting motor threshold was 49 (± 12) maximum stimulator output in the real cTBS condition and 49 (± 12) in the sham condition. The stimulation intensity used over the DLPFC (70% RMT adjusted for scalp-to-cortex distance) was 34 (± 8) maximum stimulator output in the real cTBS condition and 34 (± 8) in the sham condition. One participant reported a headache, while two participants reported discomfort following real cTBS.

### 3.2 GABA concentration

Following quality control, 4 participants with FWHM of GABA peak > 10 Hz, and 5 participants with a negative value for spectra area in at least one of the recording sessions were removed from the analysis, leaving 8 participants in the main analysis. Figure 2 shows the individual (black) and mean (red) spectra at each voxel before and after both real and sham cTBS.

There were no main effects of TIME (F_1,56_=0.13, *P*=0.72; BF_10_=0.33), CONDITION (F_1,56_=2.01, *P*=0.15; BF_10_=0.47) or SITE (F_1,56_=0.13, *P*=0.72; BF_10_=0.44) on GABA concentration and no interactions (F_1,56_=0.19, *P*=0.66; BF_10_=0.07), suggesting cTBS did not alter GABA concentration either at the stimulation site or a remote brain region (fig. 3).

**Figure 3.**
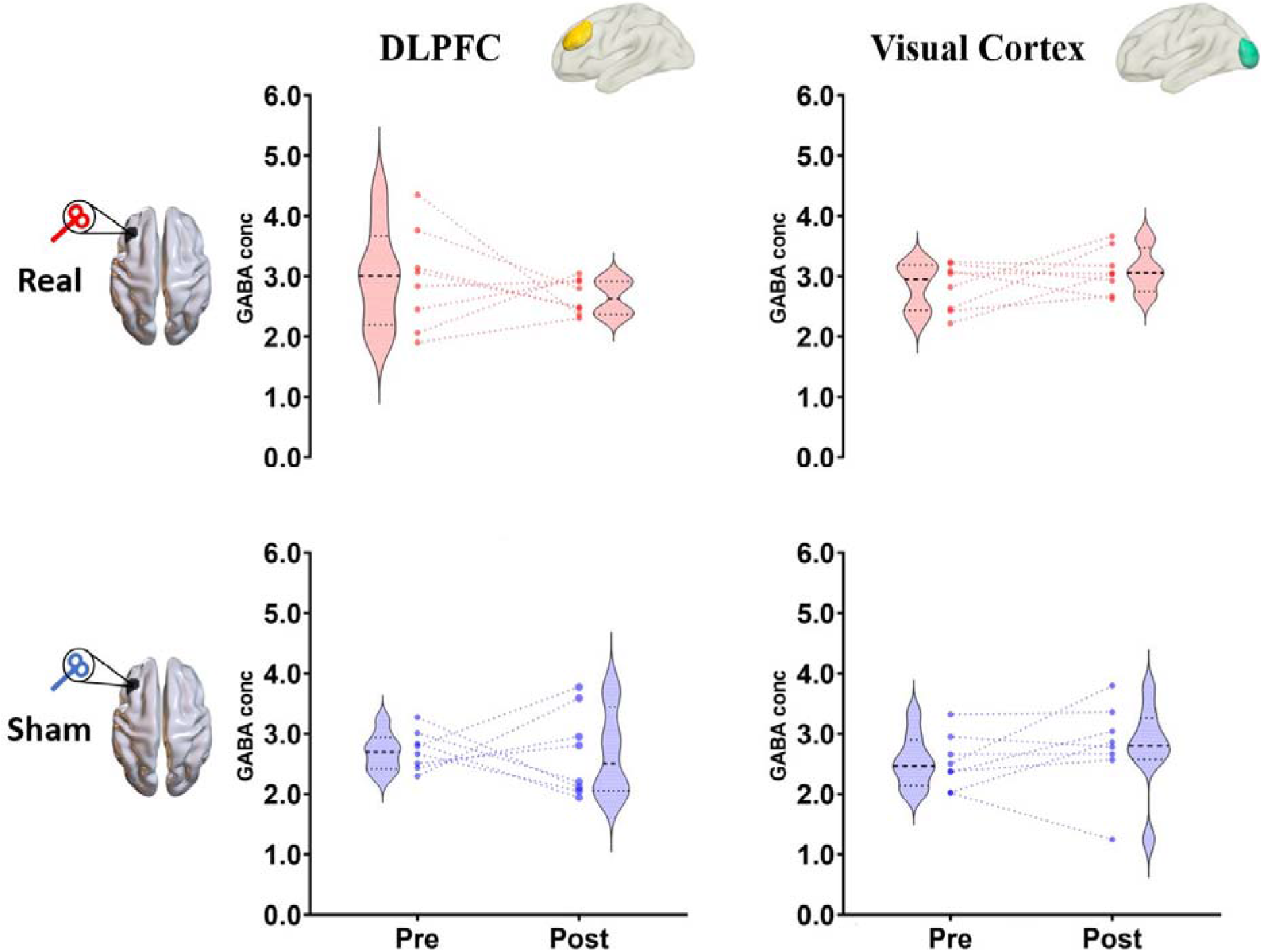
Changes in gamma aminobutyric acid (GABA) concentrations following continuous theta burst stimulation (cTBS). GABA concentrations measured from dorsolateral prefrontal cortex (DLPFC) and visual cortex voxels using magnetic resonance spectroscopy before and after real (upper panel) and sham (lower panel) cTBS. Dots and lines represent values for individual participants, whereas violin plots represent the distribution of the group data.

### 3.3 Resting state functional connectivity

Figure 4 shows seed-based resting-state connectivity maps on the cortical surface before and after real and sham cTBS. For seed-based connectivity, no significant clusters or interactions were observed using either the full-factorial design, or one-sampled t-test (all *P*>0.05, FWE-corrected), suggesting no changes in connectivity within the targeted DLPFC-network following cTBS. Furthermore, there were no main effects or interactions of GROUP or TIME when using the network-based statistic approach at any of the thresholds tested (all *P*>0.05), suggesting cTBS did not alter non-targeted networks either.

**Figure 4.**
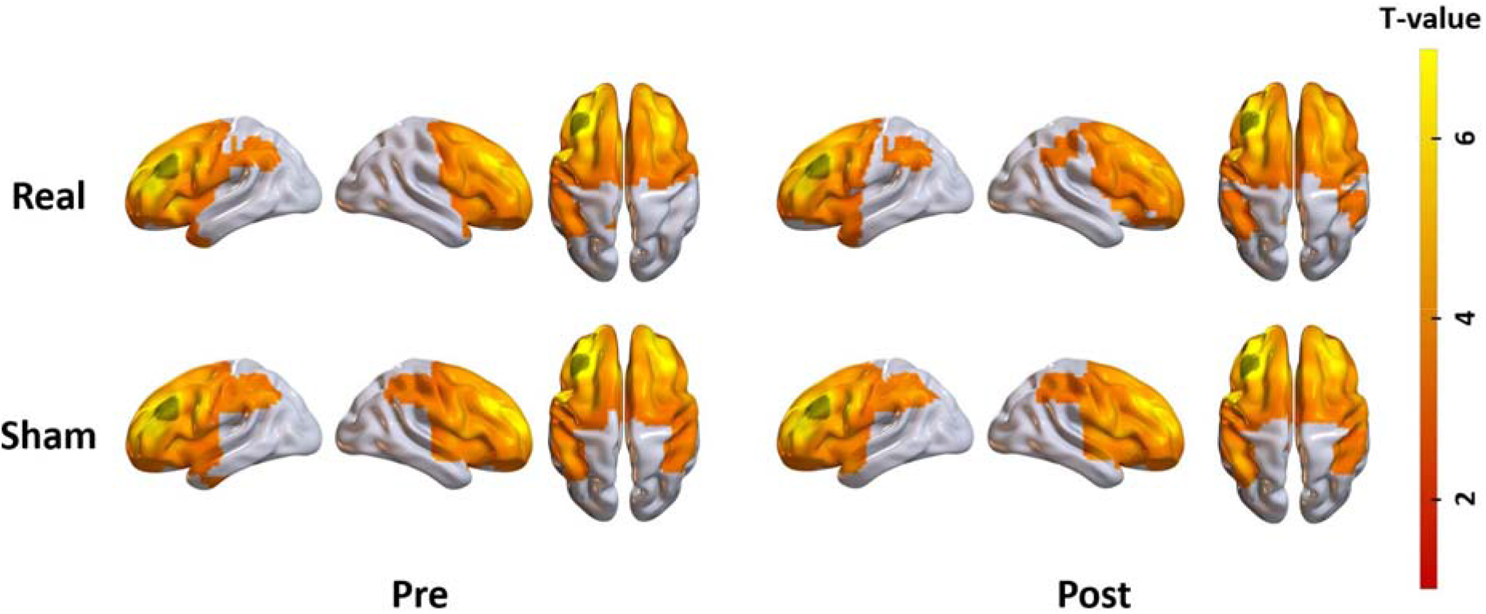
Seed-based resting-state functional connectivity maps for real and sham cTBS conditions at pre and post time points. Data have been translated to the cortical surface for visualisation and are thresholded at *P*<0.001 (one-sampled t-test). Black regions indicate the left dorsolateral prefrontal cortex stimulation site used for the seed.

### 3.4 fALFF

Figure 5 shows changes in fALFF values following real and sham cTBS. There were no main effects of TIME (F_1,64_=0.01, *P*=0.93; BF_10_=0.27) or CONDITION (F_1,64_=0.54, *P*=0.47; BF_10_=0.52) on fALFF and no interactions (F_1,64_=0.02, *P*=0.88; BF_10_=0.04), suggesting cTBS did not alter low-frequency fluctuations at the site of stimulation.

**Figure 5.**
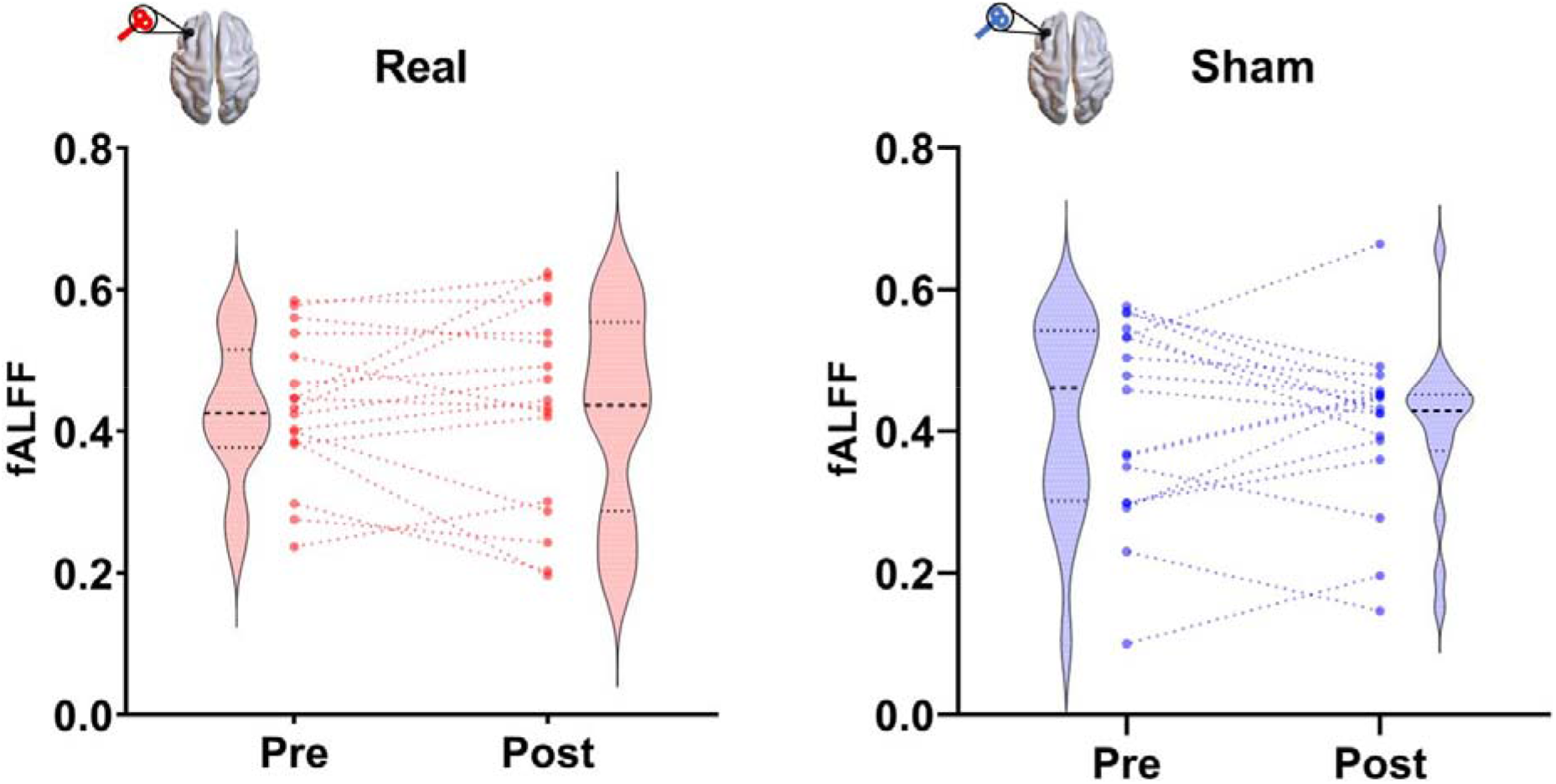
Changes in low frequency fluctuations in the dorsolateral prefrontal cortex (DLPFC) following continuous theta burst stimulation. Fractional amplitude of low-frequency fluctuation (fALFF) values measured from the time series extracted from the DLPFC seed region at the site of stimulation. Dots and lines represent individual participants, whereas violin plots represent the distribution of the group data.

### 3.5 BOLD activity during the n-back task

Figure 6 shows BOLD activation and deactivation patterns for the real and sham cTBS conditions at pre and post time points. No significant clusters were observed for any contrasts between real and sham cTBS conditions or time-points for either the full-factorial model or the one-sampled t-tests, nor were there any interactions (all *P*>0.05, FWE-corrected), suggesting cTBS to DLPFC did not alter BOLD activation patterns during the n-back task.

**Figure 6.**
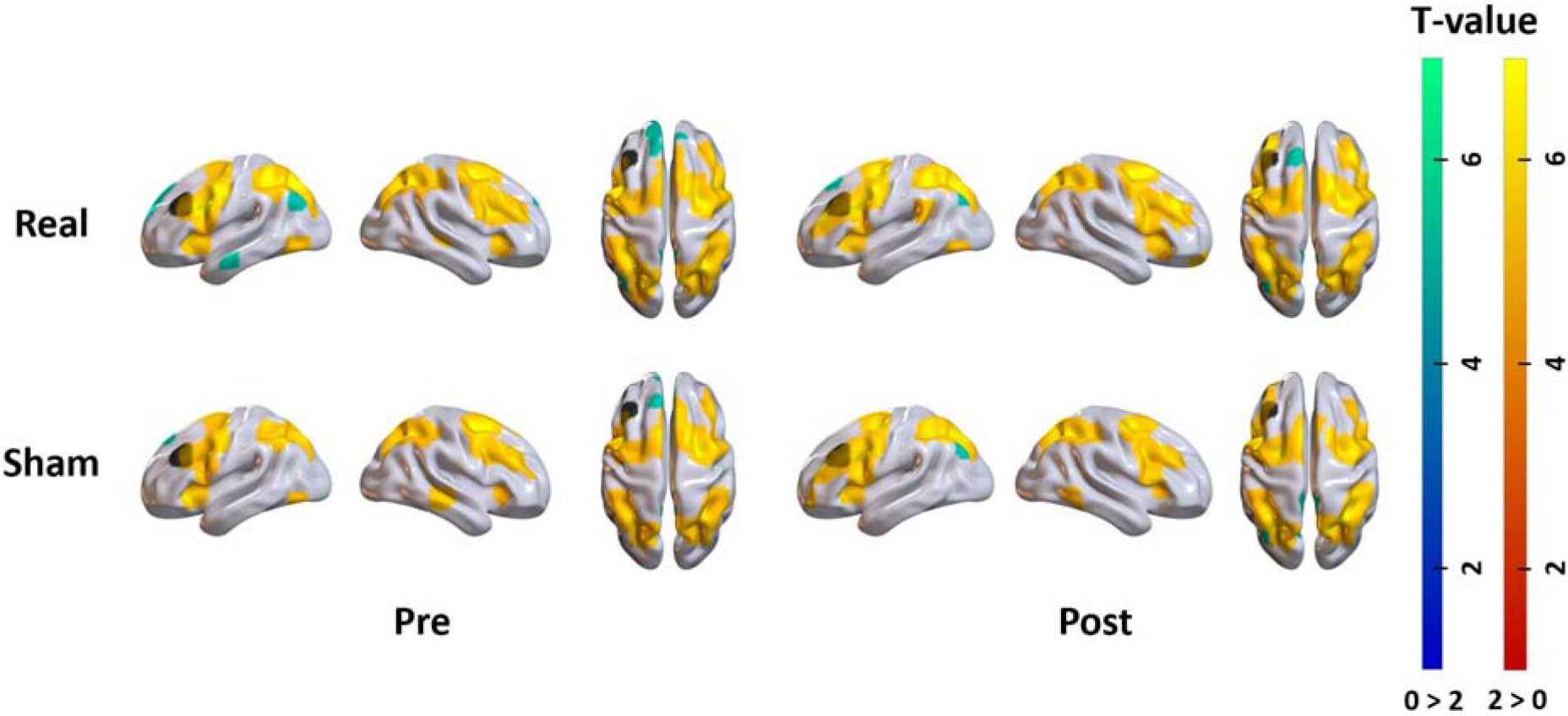
Task activation maps before and after continuous theta burst stimulation (cTBS). Blood-oxygen level dependent activation (2 back > 0 back) and deactivation (0 back > 2 back) maps during the n-back task before and after real and sham cTBS conditions. Data have been translated to the cortical surface for visualisation and are thresholded at *P*<0.001 (one-sampled t-test, thresholded at *P*<0.00001, k>10). Black regions indicate the left dorsolateral prefrontal cortex stimulation site.

### 3.6 Working memory performance

Figure 7 shows changes in *d*^*’*^ scores following cTBS. There were no main effects of TIME (F_1,63_=2.90, *P*=0.11; BF_10_=1.00) or CONDITION (F_1,63_=1.30, *P*=0.30; BF_10_=0.42) on *d*^*’*^ scores and no interactions (F_1,63_=2.60, *P*=0.13; BF_10_=0.33), suggesting cTBS over DLPFC did not alter working memory performance. Furthermore, there were no main effects of TIME (F_1,63_=4.52, *P*=0.06; BF_10_= 0.60) or CONDITION (F_1,63_=0.40, *P*=0.54; BF_10_=0.31) on RT and no interactions (F_1,63_=1.10, *P*=0.32; BF_10_=0.10), suggesting cTBS over DLPFC did not alter working memory performance.

**Figure 7.**
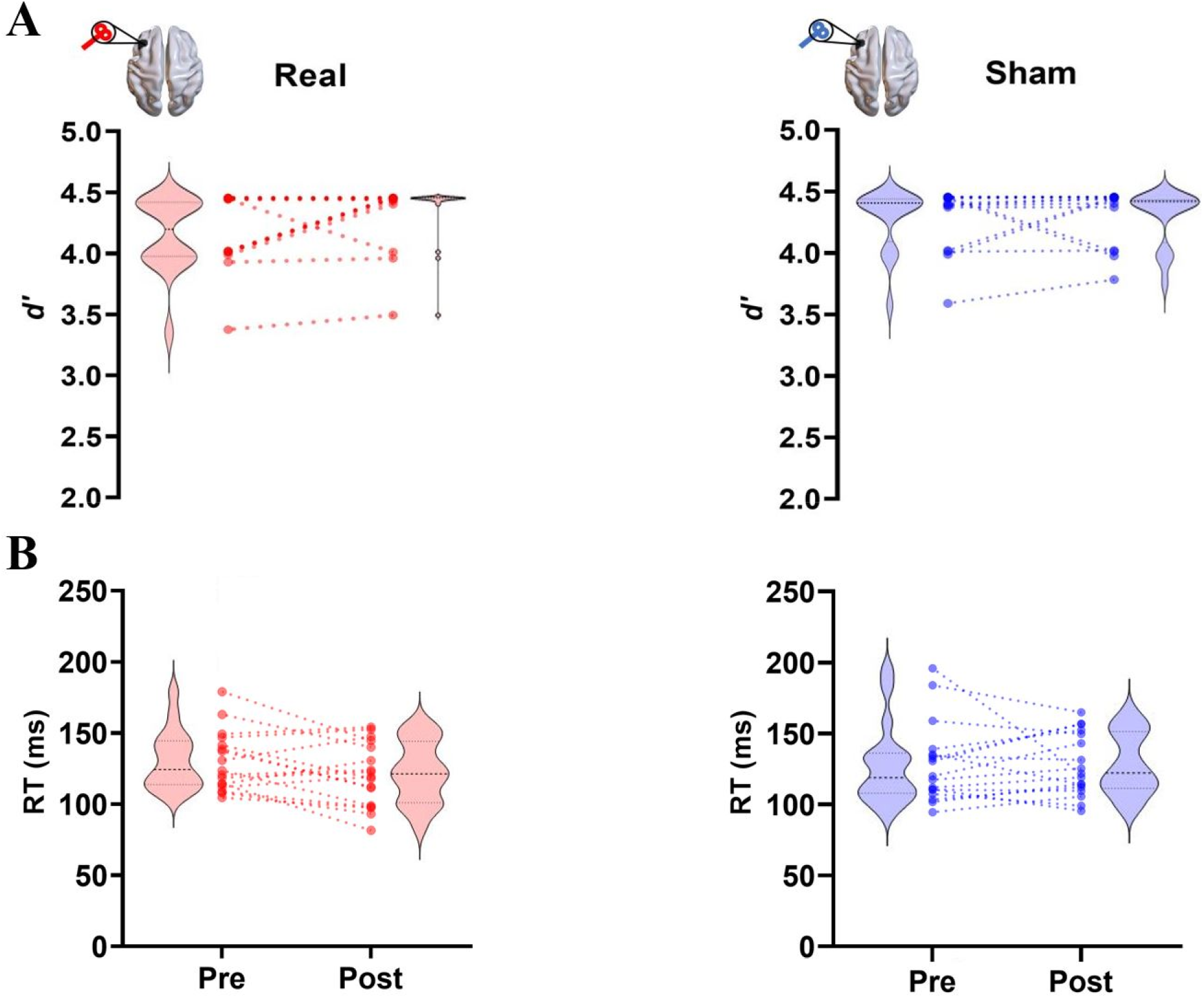
Changes in working memory performance following continuous theta burst stimulation (cTBS). *d’* scores (A) and reaction times (RT; B) measured from the 2-back task for real (red) and sham (blue) cTBS conditions at pre and post time points. Dots and lines represent individual participants, violin plots represent the distribution of the group data. Note that darker dots represent overlapping participants.

## 4. Discussion

In this study, we conducted a comprehensive, multi-modal assessment of how cTBS over DLPFC influences brain activity and working memory performance. Using MRS and fMRI, we investigated changes to GABA concentrations and low-frequency fluctuations at the site of stimulation, as well as alterations in resting-state network connectivity, and BOLD activation patterns during a working memory n-back task. We also assessed changes in performance on the 2-back task following stimulation. In contrast to our hypotheses, we could not find evidence for changes in any of these measures following cTBS over DLPFC. Where possible, these findings were complemented with Bayesian statistics, which provided consistent and strong evidence in favour of the null hypothesis. Overall, our results suggest that cTBS over DLPFC does not consistently alter brain activity patterns at a local or network-level, nor does it alter WM performance. These findings suggest that the effects of cTBS over DLPFC do not consistently alter brain activity at the group level and challenge the notion of cTBS as a universal ‘inhibitory’ paradigm.

Contrary to our hypothesis, we found no evidence of altered GABA concentration at local or remote sites following cTBS to DLPFC. Qualitatively, there was some indication of reduced variability in GABA concentration within DLPFC following cTBS, though these effects were not statistically significant. These findings contrast with past research reporting altered GABAergic transmission following application of cTBS over motor cortex (Huang et al., 2005; Murakami et al., 2008; McAllister et al., 2009; Stagg et al., 2009; Goldsworthy et al., 2013). For example, previous studies have reported increases in GABA concentration measured with MRS (Stagg et al., 2009) and reductions in TMS-based measures of cortical inhibition (Huang et al., 2005; Murakami et al., 2008; McAllister et al., 2009; Goldsworthy et al., 2013) following cTBS to primary motor cortex. Hence, our results suggest that the interaction between cTBS and inhibitory circuits observed in the motor cortex may not translate directly to non-motor brain regions such as the DLPFC. Notably, however, several past studies have identified changes in GABA concentration in non-motor regions following TBS (Cuypers & Marsman, 2021). Increases in GABA concentration have been observed in the visual cortex following cTBS (Stoby et al., 2019), and in dorsomedial prefrontal cortex following intermittent TBS to a functionally connected region of parietal cortex (Vidal-Piñeiro et al., 2015). Considered in the context of the present study, these outcomes indicate that the effects of cTBS on GABA concentration may be regionally specific, with considerable variability observed across the distinct cortical regions targeted in past studies. This perspective is also supported by a recent study which reported differential effects of multi-day rTMS on GABA concentration between parietal cortex and pre-supplementary motor area targets (Hendrikse et al., 2020b). Overall, our findings challenge the consistency of cTBS-induced effects on GABA outside of the motor system. Further research using within-subject study designs is required to identify the specific factors contributing to the variable effects of cTBS on GABA concentration across different cortical regions.

Several studies have reported changes in functional connectivity following cTBS to both motor and non-motor regions (Beynel et al., 2020). For example, a recent study reported that cTBS over DLPFC reduced functional connectivity between the stimulation target and regions of the default-mode network (Shang et al., 2020), whereas another reported reductions in motor network connectivity during a motor learning task following cTBS over primary motor cortex (Steel et al., 2016). However, the direction of connectivity changes is not always consistent, with another study reporting increases in resting-state connectivity within the motor network following cTBS (Cocchi et al., 2015). Furthermore, not all studies have reported changes in functional connectivity following cTBS, with a recent study reporting no connectivity changes following stimulation of either right superior parietal lobule or temporo-parietal junction relative to a vertex control site (Machner et al., 2021). In contrast, we found no evidence of altered functional connectivity following cTBS over DLPFC either within the stimulated network, or in non-targeted networks.

The cause of these inconsistent outcomes between studies is unclear. One possibility is that our null findings might reflect the high inter-individual variability observed following rTMS paradigms (Hamada et al., 2013; López-Alonso et al., 2014). For example, Hamada and colleagues (2013) found that only ∼50% of individuals showed a reduction in MEP amplitude when cTBS was applied to primary motor cortex in a large sample (n=56), resulting in no significant changes in MEP amplitude at the group level. Alternatively, the lack of changes in functional connectivity might reflect a site-specific property of the DLPFC. Site-specific changes in functional connectivity have been reported following rTMS (Cocchi et al., 2016; Castrillon et al., 2020; Hendrikse et al., 2020a), with certain studies reporting divergent changes in connectivity when identical forms of stimulation were applied across different cortical regions (Cocchi et al., 2016; Castrillon et al., 2020). Overall, this evidence suggests that the effects of rTMS on functional connectivity are highly variable and may not follow frequency-dependent conventions (e.g., reduced connectivity following 1 Hz rTMS and cTBS), particularly when stimulation is applied to non-motor regions.

The DLPFC is a primary node in the network of brain regions supporting working memory function (Rottschy et al., 2012). Hence, in this study, we hypothesised that cTBS over DLPFC would impair performance on a working memory task (n-back). Contrary to our hypotheses, we did not observe any evidence of changes in working memory performance working memory performance (d’ scores or RT), nor altered BOLD activity within DLPFC or other regions supporting working memory. These results conflict with the findings of two previous studies reporting reductions in n-back performance following cTBS over DLPFC (Schicktanz et al., 2015; Vékony et al., 2018), but are in-line with another study reporting no effect of cTBS on working memory performance (Viejo-Sobera et al., 2017). Given that cTBS did not alter GABA concentrations or resting DLPFC activity/connectivity in our sample, it is perhaps not surprising that we did not observe corresponding change in working memory performance or activity patterns. To more reliably study the relationship between DLPFC activity/connectivity and working memory, future research should focus on developing rTMS paradigms capable of consistently altering brain activity in regions like the DLPFC.

There are several limitations of this study that should be acknowledged. First, our analyses were conducted on a relatively small sample of 17 healthy adults. Our sample was further reduced to 8 participants for GABA concentration measures due to our conservative quality control approach, a step which is required to ensure robust conclusions when using MRS (Alger, 2010). To our knowledge, this study is the first to provide a comprehensive multi-modal assessment of the effects of cTBS over DLPFC, and thus represents the largest study of its kind to-date. However, considering the variability associated with cTBS (Hamada et al., 2013), this sample size may have lacked the required experimental power to detect small/moderate effects. While the combined use of frequentist and Bayesian statistics provided consistent, strong evidence in support of the null hypothesis, future studies using larger sample sizes are required to establish the reliability of these findings. Second, while stereotactic neuronavigation was used to account for individual differences in neuroanatomy, we derived our DLPFC target from standardized MNI coordinates [x = - 38, y = 30, z = 30] – an approach commonly adopted in the field. In both experimental and clinical contexts, there is evidence to suggest that personalising stimulation targets on the basis of individual functional connectivity maps may reduce response variability (Sack et al., 2009; Cash et al., 2020; Cash et al., 2021). Therefore, tailoring stimulation to subject-specific regions of DLPFC with strongest connectivity to task-relevant functional networks may have induced more consistent effects on both cognition and connectivity. However, in this study, we observed no change in local neural activity or GABA concentration, suggesting that cTBS may not have induced the significant local effects on DLPFC activity to influence connectivity across a wider network-scale. Further, personalised methods may not always enhance the reproducibility of rTMS effects (Hendrikse et al., 2020a). Finally, our working memory assessments occurred approximately 30-40 mins after stimulation. While early studies suggested cTBS after-effects lasted up to 60 mins (measured by reductions in MEPs; (Huang et al., 2005)), more recent meta-analyses have suggested these effects are strongest within the first 20-30 mins (Chung et al., 2016). As such, our design may not have been optimal for detecting changes in working memory performance and task-related brain activity.

## 5. Conclusion

In summary, we found no evidence that cTBS over DLPFC modulates GABA concentration, functional connectivity or working memory performance using a comprehensive, multi-modal assessment of brain activity. Our findings add to a growing body of literature questioning the reliability of cTBS for altering brain activity in motor and non-motor regions (Huang et al., 2017). The development and validation of more reliable and consistent rTMS protocols is required to better study the causal relationship between DLPFC function and working memory.

## 6. Acknowledgements

We thank Dr. Ian Harding and Prof. Nellie Georgiou-Karistianis for sharing the code for the n-back task, Dr. James Coxon for sharing the code to generate figures for MR spectroscopy, and the staff of Monash Biomedical Imaging for assistance in designing the MRI sequences and collecting the data.

## 8. Data and Material Availability

Readers seeking access to the data should contact Tribikram Thapa. Access will be granted to named individuals in accordance with ethical procedures governing the reuse of sensitive data. Specifically, requestors must complete a formal data sharing agreement and make clear the process by which participant consent would be sought and data used. Deidentified behavioural data and inputs for group level fMRIanalyses (i.e., 1^st^ level fMRI contrast images) are available at https://osf.io/x467g/, and all code used for analyses is available at https://github.com/TribThapa/cTBS_Study.

